# Enhancing Quantitative and Data Science Education for Graduate Students in Biomedical Science

**DOI:** 10.1101/2021.12.03.471108

**Authors:** Louis J. Gross, Rachel Patton McCord, Sondra LoRe, Vitaly V. Ganusov, Tian Hong, W. Christopher Strickland, David Talmy, Albrecht G. von Arnim, Greg Wiggins

## Abstract

Substantial guidance is available on undergraduate quantitative training for biologists, including reports focused on biomedical science, but far less attention has been paid to the graduate curriculum. In this setting, we propose an innovative approach to quantitative education that goes beyond recommendations of a course or set of courses or activities. Due to the diversity of quantitative methods, it is infeasible to expect that biomedical PhD students can be exposed to more than a minority of the quantitative concepts and techniques employed in modern biology. We developed a novel prioritization approach in which we mined and analyzed quantitative concepts and skills from publications that faculty in relevant units deemed central to the scientific comprehension of their field. The analysis provides a prioritization of quantitative skills and concepts and could represent an effective method to drive curricular focus based upon program-specific faculty input for biological science programs of all types. Our results highlight the disconnect between typical undergraduate quantitative education for life science students, focused on continuous mathematics, and the concepts and skills in graphics, statistics, and discrete mathematics that arise from priorities established by biomedical science faculty.

**One Sentence Summary:** We developed a novel approach to prioritize quantitative concepts and methods for inclusion in a graduate biomedical science curriculum based upon approaches included in faculty-identified key publications.

## Introduction

There is widespread agreement that training in quantitative approaches is critical for graduate students in biomedical fields (*1*). Less clarity exists about how to prioritize which quantitative concepts and approaches are most important for students to learn. In general, there has been more explicit study of how to incorporate quantitative skills into undergraduate biology programs than there has been at the graduate level (*2-5*). There are hosts of different quantitative topics, conceptual approaches, and skills, and so training typically involves trade-offs on the concepts taught. General guidance for curricular development is provided by reports of authoritative groups in professional societies, accreditation agencies, and committees supported by federal agencies and foundations through organizations such as the National Academies of Science, Engineering and Medicine. Aside from these reports, previous efforts to identify key quantitative concepts for students in a field have typically taken two forms: (i) automated text mining of publications, or (ii) faculty deliberation and decision-making committees. For example, large samples of articles from field-specific journals have been mined for pre-defined statistical terms to inform which concepts are key to graduate training in higher education research (*6, 7*), ecology (*8*), or oncology (*9*). In the second approach, there are examples from geoscience departments (*10*), life science programs (*11*), and business schools (*12*) which used faculty meetings or surveys of faculty, graduates, or employers to explicitly define key quantitative skills deemed essential for students in these fields. We propose an alternative approach to guide the enhancement of quantitative components of the PhD curriculum based upon local needs as suggested by faculty in relevant units. We asked faculty from three programs that train biomedical science graduate students to identify papers that are key to understanding their field. We then mined the papers through expert coding to obtain the quantitative approaches represented. This approach allows us to gain the benefit of both broad-based faculty input and supervised data filtering. These data were analyzed and compared to information on the prior educational background and syllabi of courses taken by students in these programs.

## Materials and Methods

### Data collection methodology

For the first phase, qualitative data was collected in the form of document analysis of research articles from the faculty associated with the major programs at the University of Tennessee, Knoxville (UTK) which educate graduate students in biomedical science - the Departments of Biochemistry & Cellular and Molecular Biology (BCMB), Microbiology (Micro), and the University of Tennessee□Oak Ridge National Lab (UT-ORNL) Graduate School of Genome Science & Technology (GST). During the semester of Fall 2018, faculty associated with these programs were asked to provide a single journal article published in the last five years which they considered important for all the students in their graduate program, not just those associated with their lab, to be able to read with comprehension. These articles may be ones used in their courses and seminars, but this was not emphasized. Faculty were asked not to submit review articles and were not told to emphasize quantitative topics in the papers suggested, but to submit papers with important scientific content. Solicitations were conducted over seven weeks, resulting in 48 papers submitted from 40 respondents. The faculty came from three core graduate programs: BCMB, Micro, and GST and a few faculty with main appointments in additional units (Biomedical and Diagnostics, Nutrition, UT Medical Center, Plant Sciences, Biosystems Engineering and Soil Science, Animal Science, and Electrical Engineering and Computer Science) were also involved because of their affiliation with the core programs (Table S1).

The research methods employed an exploratory sequential mixed methods design (*13, 14*) where qualitative methods are followed by quantitative measures (Figure 1). After the collection of articles, the six faculty on the project were asked to identify quantitative skills from a sampling of eight randomly assigned papers spending 10-15 minutes identifying the quantitative tools/skills and/or concepts used in each article. Meeting facilitators and the evaluator, acting as a participant-observer (*15, 16*), kept ethnographic memos (*17*) regarding quantitative concepts and skills discussed. The initial review of article samples concluded with a listing of 173 quantitative skills under 21 general concepts. After consideration of overlaps in the quantitative topics and general concepts, this list was distilled to an aggregated listing of seven general concepts and 68 specific skills associated with the general concepts.

**Figure 1.**
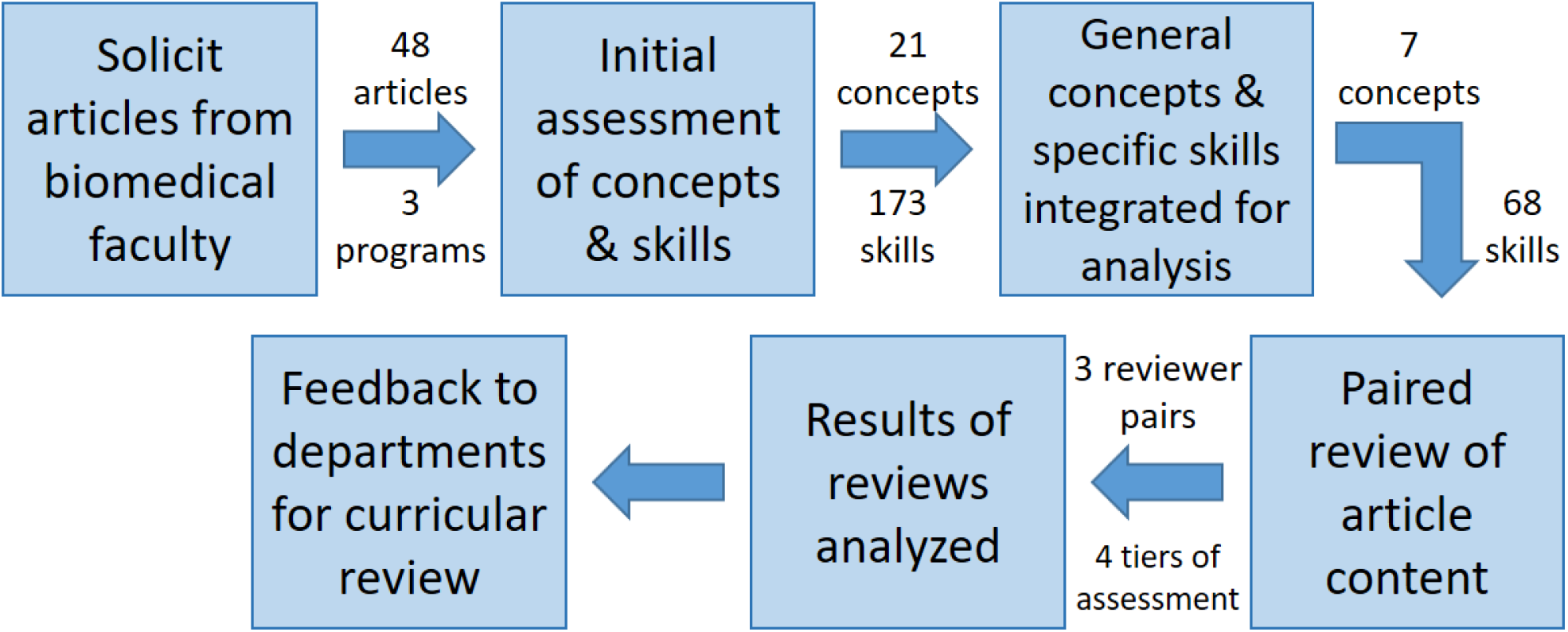
Process for data collection and analysis of articles chosen by faculty as appropriate for all biomedical students completing a PhD in their program to read with comprehension. See main text for details of the steps taken for collection of data and analysis.

The quantitative phase of the mixed methods research design included the creation of a survey to be used to re□analyze the submitted papers. The articles were then grouped with teams of two members of the faculty team assigned to consider in detail each of a set of papers that were deemed to be most connected to their backgrounds. The second analysis of the journal article’s quantitative content included four tiers of related assignments for each concept/skill: 1) the presence of generalized concepts in the sample paper, 2) the level of importance of this concept to understanding the paper, followed by 3) specific skills related to the general concept and 4) the level of importance of the specific skill to understanding the paper. The rankings for each concept or skill assessed in these tiers was: 1: not present, 2: marginally important to comprehension of the paper, 3: somewhat important to comprehension of the article, or 4: very important to comprehension of the paper (Figures 1&2).

**Figure 2.**
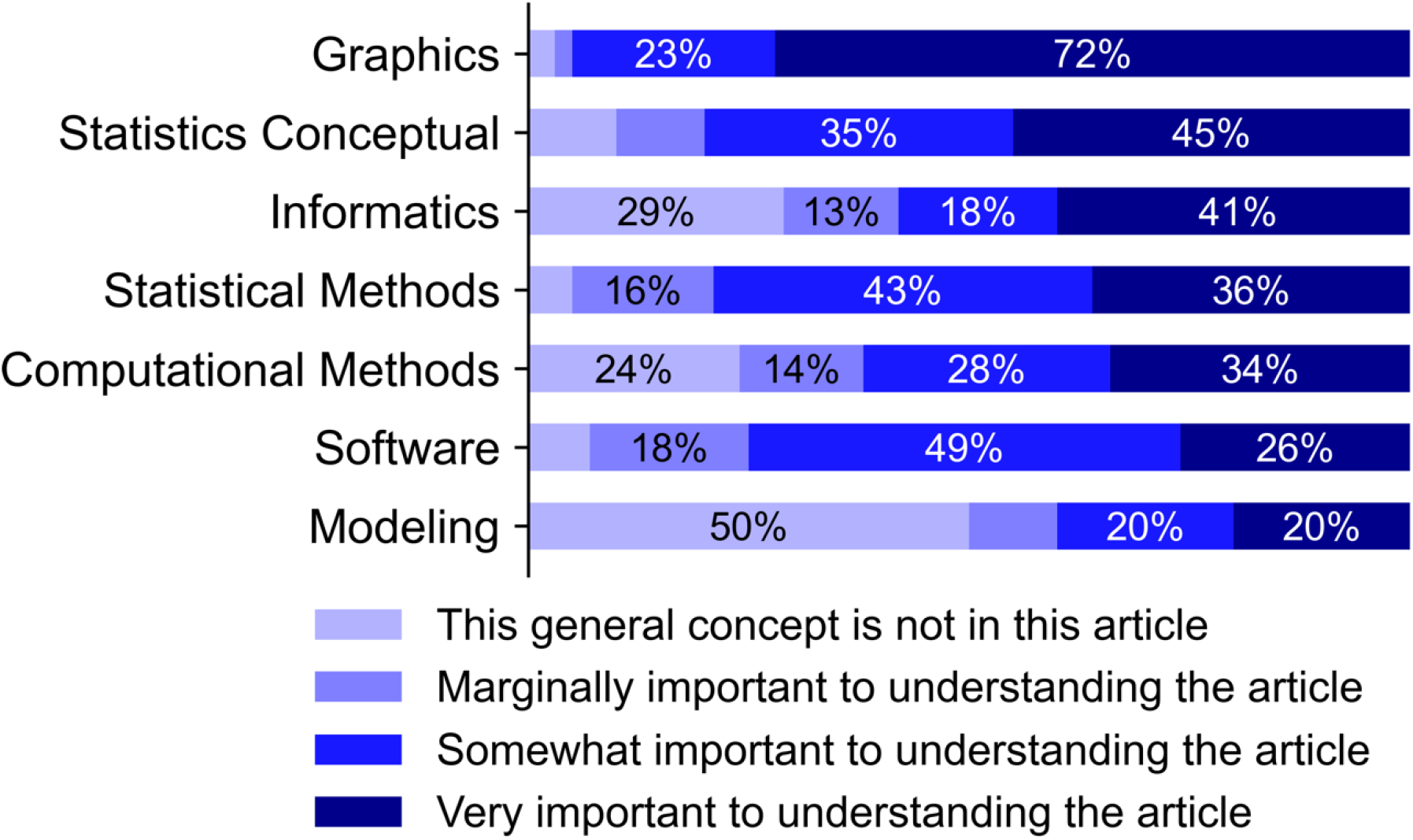
The seven general concepts identified in the submitted papers and the proportions of importance level of each concept. Percentage values indicate the percent of papers in which a given concept was assigned a given importance level. Values greater than 10% are labeled with text.

This second analysis was carried out by teams of two faculty members, who each provided their own ranking of importance of the concepts and skills. After each review pair completed the surveys for their papers the evaluator compared the results by paper. All of the paper scores were averaged collectively by concepts and specific skills. Graphics were constructed to show the existence and importance of the concepts and presented to the research team member.

### Statistical data analysis

To visualize the fractions of importance levels (1: ‘not present’, 2: ‘marginally important’, 3: ‘somewhat important’, and 4: ‘very important’) for all seven general concepts, first the overall fractions of the four levels across all concepts were calculated (Fig. S1, dashed lines). These overall fractions are defined as ‘expected’ fractions. Next, the fractions of the four levels for individual concepts were calculated (Figure 2 and Figure S1, grey bars), and plotted to visualize the deviation of the individual fractions from the expected fraction (Figure S1). For example, the fraction of ‘not present’ level for the ‘Modeling’ concept was higher than expected by 0.31, and the fraction of ‘very important’ level for the same concept was lower than expected by 0.19. For each concept, all 48 papers (i.e., 96 evaluations) were included for calculating the fractions. A chi-squared test was performed for each concept with the frequencies of the four levels and the expected frequencies derived from the overall fractions (levels were treated as categorical rather than ordinal data).

A similar method was used to visualize the importance levels of skills. For each general concept first, the overall (i.e. expected) fractions of the four levels among the evaluations in which the concept is present were calculated (Figure 3 grey bars). Then the fractions of the four levels for individual skills were calculated. A heatmap was used for each concept to visualize the deviation of the individual fractions (per skill per level) from the expected ones (Figure 3). The fraction per skill per level was with respect to evaluations in which the concept is present, except for a few instances (<3% among the 96 evaluations) where the skill levels were missing and the fractions were with respect to available evaluations. In the heatmap, white color means expected fraction, red color means higher-than-expected fraction, and blue color means lower-than-expected fraction.

**Figure 3.**
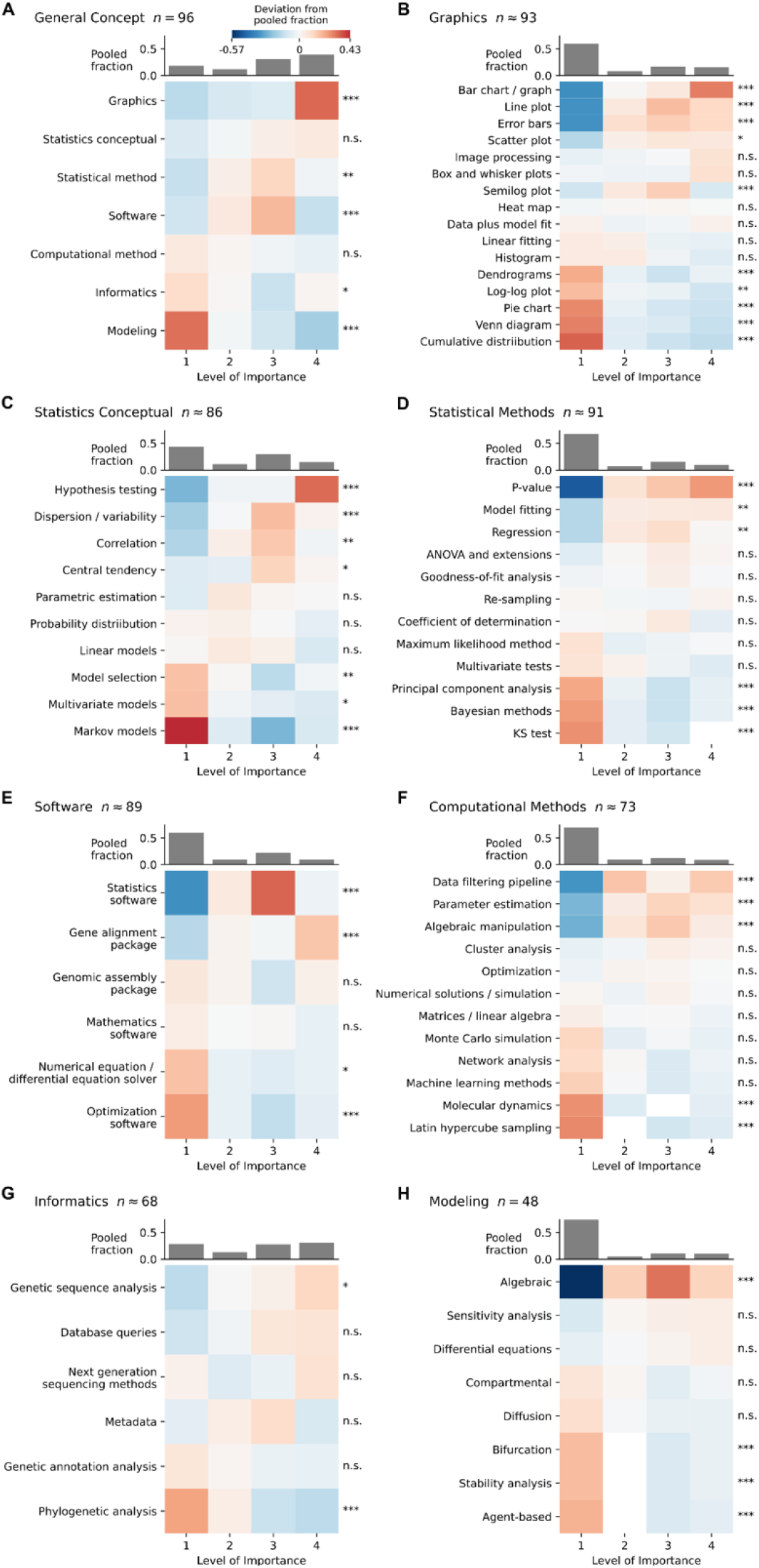
Deviations from average importance distributions for concepts (A) and skills (B to H) identified from solicited papers. Importance ranges from 1= Concept not in this article to 4 = Very important to understanding this article, as in Fig. 2. Significant deviation from expected fraction is as follows: n.s. *p* > 0.05; ^*^ *p* ≤ 0.05; ^**^ *p* ≤ 0.01; ^***^ *p* ≤ 0.001. Further detail about the data is provided in the supplementary material.

### Power analyses

To evaluate how many papers contain a given number of concepts or skills power analyses were performed. For the concept analysis, all evaluations were resampled with replacement 1,000 times and counted the percent of times whether a given concept (out of a set of seven) was present in the sample at a particular level (defined by a cut-off). For example, cut-off of 3 implied that this specific concept was somewhat important to understand the paper.

Analysis suggests that 5 papers already have most of the concepts represented (Figure S2A). A similar power analysis for the skills when resampling all evaluations suggests that different numbers of papers were needed to cover skills in different concepts (Figure S2B). For example, Informatics and Statistics: Conceptual skills were most contained in most papers while Computational Methods and Modeling required a larger sample size (20-25) to have their skills well represented. This analysis provides evidence that there was a sufficient sample size of articles to identify most of the defined skills which perhaps is not surprising given that these concepts were mined from the papers.

## Results

This case study focuses on the major units at the University of Tennessee, Knoxville (UTK) which educate PhD students in biomedical science: the Departments of Biochemistry & Cellular and Molecular Biology (BCMB), Microbiology (Micro), the University of Tennessee-Oak Ridge National Lab (UT□ORNL) Graduate School of Genome Science & Technology (GST). The three graduate programs at the core of the analysis carried out here host a total of ∼160 graduate students, predominantly in the PhD track. Microbiology includes two main areas: environmental microbiology and microbial pathogenesis. BCMB encompasses mainly three areas: physical biochemistry, and molecular and cellular biology of both plant and animal systems. GST is an intercollegiate life science program that emphasizes training across the interface of the wet-lab and dry-lab, e.g., in computational biology (*18*). It includes faculty members from Microbiology, BCMB, and other departments across the university.

Faculty associated with these units were asked to each identify at least one recently published journal article, not necessarily with quantitative emphasis, that they suggest biomedical science students completing a PhD in their program should be able to read with comprehension (Figure 1). The project team then analyzed these papers for quantitative concepts and methods, grouping these in a hierarchical manner based on a few core concepts, approaches, and skills necessary to adequately understand faculty-selected papers (see Materials and Methods for the full design description).

We obtained the distributions of the evaluations over the four importance levels (Figure 2 and Figure S1) and found that five out of the seven general concepts showed significant deviations from the pooled distributions across concepts suggesting that some quantitative concepts are more important than others for biomedical graduate students in the UT programs as judged by the faculty in those programs. Interestingly, among the seven general concepts, ‘Graphics’ was ranked at the top in terms of the median importance levels, whereas ‘Modeling’ was considered the least important concept as it was absent from most of the articles (Figure 2 and Figure S1). In addition, ‘Software’ and ‘Statistical Methods’ showed a significant deviation because they appeared in a large number of articles with a medium level of importance. Within individual concepts, 54% of the specific skills showed significant deviations in importance from the pooled distributions across all skills (Figure 3). Among skills that were considered most important, ‘Bar chart/graph’, ‘Line plot’, ‘Error bars’, ‘Hypothesis testing’ and ‘P-value’ were not only present in most of the articles that contain the corresponding general concepts, but also scored as ‘very important’ in those articles. Some skills were considered significantly important even though their general concepts did not rank high, including ‘Statistics software’, ‘Gene alignment package’, ‘Gene sequence alignment’, ‘Data filtering pipeline’, ‘Algebraic manipulation’ and ‘Parameter estimation.’ Most of these skills showed medium levels of importance in the articles where the skills were present.

Having identified which concepts and skills were noteworthy in the papers important to faculty training biomedical graduate students, we considered how these skills align with typical training of incoming graduate students in these programs. We are unaware of any formal analysis of the quantitative background of students entering US life science graduate programs. Informal evidence from students in our life science departments indicates that there is considerable variability both within programs and between programs of the formal mathematics, statistics, and computing backgrounds of these students. Curriculum requirements for undergraduate biology programs, including that at UTK, typically include calculus, but not necessarily statistics and/or data analysis/visualization courses or any specific requirements in computational biology (*19, 20*). This suggests that many of the mathematical skills needed to engage with topics deemed important in our current study will need to be built during the graduate biomedical training process.

## Discussion

This study developed a novel and generalizable data collection methodology targeting faculty opinions about core concepts for their PhD students in the biomedical sciences. Upon analyzing the supplied manuscripts for their quantitative, data-analytical techniques our results suggest that there is significant variability across both general concepts and particular skills with regard to their estimated importance in comprehending key papers. It is important to note that scoring of the articles was primarily based on a standard of literacy; that is, the goal was to determine the subject-area knowledge necessary to *comprehend* the material in question. An independent group of researchers who are not specific, subject-matter experts in a paper’s field of study will frequently have difficulty determining the technical knowledge necessary to reproduce the study. Consequently, determining the quantitative, data-analytical techniques necessary for reproduction was beyond the scope of our work. Our comprehension-based approach has the side-effect of emphasizing quantitative techniques that have an intrinsic communication or presentation aspect (e.g., ‘Graphics’) at the expense of other categories (e.g., ‘Computational Methods’) which are more fully methods- or analysis-based. An additional limitation to our comprehension-based approach is that it is inherently biased toward subject matter whose prerequisite knowledge is ubiquitous among current and past biomedical researchers and is therefore a measure of the status-quo for biomedical research literacy rather than an indicator of field direction. More complex methods or methods requiring a high degree of uncommon background knowledge are less likely to be represented here, regardless of their merits for advancing biomedical research.

Our methodology reveals critical details about the baseline of knowledge necessary for anyone pursuing a PhD in the biomedical sciences today. It also provides a field-specific, detailed usage comparison of quantitative tools: bar and line plots, hypothesis-testing, statistical software packages, p-values, algebraic expressions, and data analysis software all stand out as highlights of research methodology while dynamic modeling and more advanced data-science techniques are more uncommon. The results indicate a potential disconnect between the typical formal quantitative training included in undergraduate life science, focused on calculus and basic statistics, and the focal concepts identified in our analysis on visualization, statistical and computational methods. Understanding and quantifying this disconnect can help to inform what quantitative concepts and skills should be emphasized in the graduate biomedical curriculum.

Our analysis suggests the types of skills that should either be already present in UTK biomedical graduate students at admission or that need to be learned in graduate school. These results have strong implications for curricula of STEM-training undergraduate programs, particularly if they prove generalizable to other graduate programs training students in biomedicine. The current strong focus on continuous mathematics (e.g., calculus I and II) may need to be replaced with other mathematics-based courses that put stronger emphasis on data analysis and interpretation. Furthermore, biomedical graduate programs may need to start offering courses in data science to align training with expectations of the faculty. We would welcome further studies examining whether our results are a part of a general pattern or an outlier among the general expectations of core concepts and skills that graduate students in biomedical sciences must have.

Our analysis suggests the types of skills that should be acquired during the course of biological/biomedical graduate work. We envision that such changes can be implemented as a three□tiered sequence of structured training elements. For entry□level PhD trainees we propose to raise ‘Awareness’ through formal learning units to be utilized in entry□level bioscience graduate courses illustrating past (and current) trends in quantitative life science. For mid□level trainees, we propose to create ‘Keys to Success’ through short and intensive training vehicles that build competence for actionable, quantitative skills based on carefully chosen experimental designs and biological data. Finally, we propose to foster a self□sustaining ‘Peer□Learning Community’ by networking advanced students (3rd year and above) with junior trainees (below 3rd year) in the form of tutorials and user groups. Together, the objective is that these training elements will change the mindset of biomedical PhD students, so they assimilate the idea that an integrated collection of quantitative skills is fundamental to their career success.

## Author contributions

Conceptualization (all authors), Data collection and curation (all authors), Formal analysis (VVG, WCS, TH), Funding acquisition (LJG), Investigation (all authors), Methodology (LJG), Project administration (LJG, GJW), Validation and visualization (VVG, TH, WCS, SL), Writing original draft (LJG, WCS, VVG, TH, RPM), Writing, reviewing and editing (all authors)

## Data and materials availability

All data is available in the main text, the supplementary materials or upon request to the authors.

