## Supplemental information for "Enhancing Quantitative and Data Science Education for Graduate Students in Biomedical Science"

**This PDF file includes:**

Figs. S1 and S2

Table S1

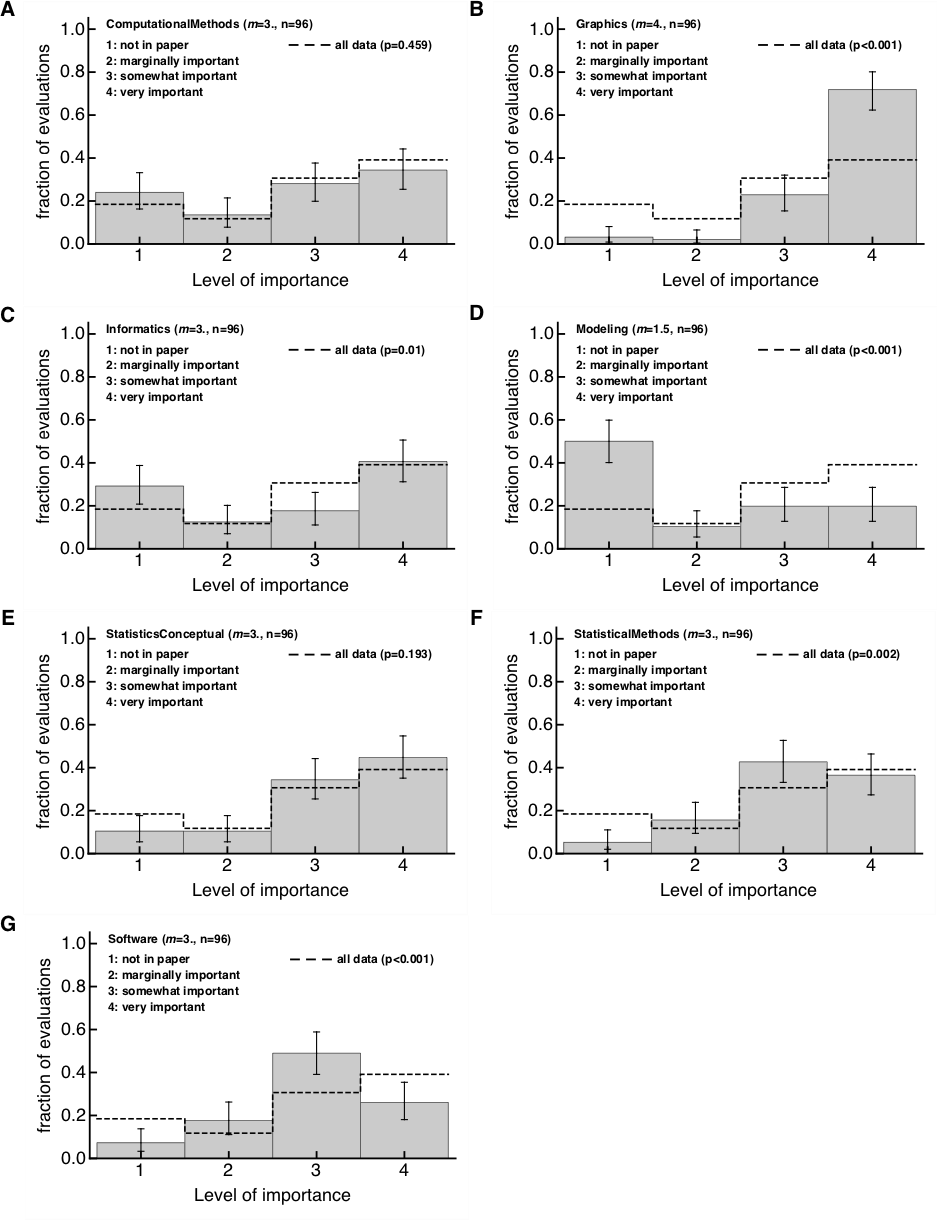

**Fig. S1. Ranking of seven different concepts by the level of importance to understand 48 scientific papers chosen by the biomedical sciences-related faculty at the University of Tennessee.** For all papers we generated 2 evaluations that ranked different concepts on the level of importance as being absent in the paper (level 1), being marginally important (2), somewhat important (3) or very important (4) for understanding of the paper. Analysis was done for Computational Methods (A), Graphics (B), Informatics (C), Modeling (D), Statistical Concepts (E), Statistical Methods (F), and Software (G). Number of evaluations per concept (*n*) and the median ranking (*m*) are indicated on individual panels. Dashed lines in each histogram represent distribution of rankings of all pooled data, and indicated p-values are from the Pearson’s chi square test for the difference between pooled data and data for the specific concept. Confidence intervals for each ranking were calculated using Jeffrey’s intervals for binomial proportions (Brown et al. Stat Sci 2001).

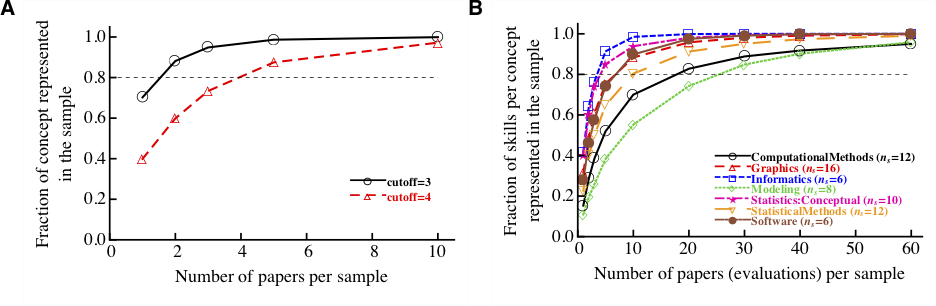

**Fig. S2. Most concepts and skills are well represented in the sample.** We performed power analysis in which we resampled evaluations and determined whether a given concept (A) or a given skill within a concept (B) is present in the sample. Simulations were done 1,000 times with replacement. Whether concept or skill was present in the sample was determined by a cut-off (A: at least at level 3 or at least at level 4, or B: level 3 or higher). Number of skills per concept in panel B (*n_s_*) for different concepts is noted for individual concepts.

| **Contributor Department** | **Paper Title** | **Initial Evaluator^1^** | **Evaluator Pair^2^** |
| --- | --- | --- | --- |
| Biochem. Cell. & Mol. Biol. | R. S. Sorenson, M. J. Deshotel, K. Johnson, F. R. Adler, L. E. Sieburth, *PNAS* **115**, E1485-E1494 (2018). | 6 | 2 & 6 |
| Biochem. Cell. & Mol. Biol. | K. Holbrook *et al., Mol. Plant* **9**, 1286-1301 (2016). | 3 | 2 & 6 |
| Biochem. Cell. & Mol. Biol. | R. Ahrends *et al., Science* **344**, 1384-1389 (2014). | 5 | 2 & 6 |
| Biochem. Cell. & Mol. Biol. | M. J. P. Chaisson *et al., Nature* **517**, 608-611 (2015). | 6 | 3 & 5 |
| Biochem. Cell. & Mol. Biol. | P. A. Ott *et al., Nature* **547**, 217-221 (2017). | 5 | 1 & 4 |
| Biochem. Cell. & Mol. Biol. | V. Nguyen *et al., Science* **355**, 289-294 (2017). | 3 | 3 & 5 |
| Biochem. Cell. & Mol. Biol. | N. A. McDonald, A. L. Lind, S. E. Smith, R. Li, K. L. Gould, *Elife* **6**, e28865 (2017). | 5 | 2 & 6 |
| Biochem. Cell. & Mol. Biol. | F. O. Bendezu *et al., PLOS Biology* **13**, e1002097 (2015). | 3 | 2 & 6 |
| Biochem. Cell. & Mol. Biol. | A. Zotter, F. Bäuerle, D. Dey, V. Kiss, G. Schreiber, *J. Biol. Chem.* **292**, 15838-15848 (2017). | 5 | 1 & 4 |
| Biochem. Cell. & Mol. Biol. | I. Yu *et al., Elife* **5**, e19274 (2016). | 1 | 3 & 5 |
| Biochem. Cell. & Mol. Biol. | G.-H. Lim *et al., Cell Host Microbe* **19**, 541-549 (2016). | 1 | 2 & 6 |
| Biochem. Cell. & Mol. Biol. | Y. Gao, W. Yang, *Science* **352**, 1334-1337 (2016). | 4 | 2 & 6 |
| Biochem. Cell. & Mol. Biol. | A. M. Valm *et al., Nature* **546**, 162-167 (2017). | 2 | 2 & 6 |
| Biochem. Cell. & Mol. Biol. | J. R. Dixon *et al., Nature* **485**, 376-380 (2012). | 4 | 3 & 5 |
| Biochem. Cell. & Mol. Biol. | C. M. Denais *et al., Science* **352**, 353-358 (2016). | 6 | 1 & 4 |
| Biochem. Cell. & Mol. Biol. | D. S. Alves *et al., Elife* **7**, e36645 (2018). | 4 | 3 & 5 |
| Biochem. Cell. & Mol. Biol. | C. Merchante *et al., Cell* **163**, 684-697 (2015). | 1 | 3 & 5 |
| Biochem. Cell. & Mol. Biol. | T. M. Lowe-Power *et al., Environ. Microbiol.* **20**, 1330-1349 (2018). | 4 | 3 & 5 |
| Biochem. Cell. & Mol. Biol. | A. Zeisel *et al., Cell* **174**, 999-1014.e22 (2018). | 1 | 3 & 5 |
| Genome Sci. Tech. | I. Johansson, A. Esberg, L. Eriksson, S. Haworth, P. Lif Holgerson, *PLOS One* **13**, e0193504 (2018). | 3 | 1 & 4 |
| Genome Sci. Tech. | B. Inceoglu *et al., PNAS* **112**, 9082-9087 (2015). | 4 | 3 & 5 |
| Genome Sci. Tech. | D. B. Richards *et al., N. Engl. J. Med.* **373**, 1106-1114 (2015). | 5 | 1 & 4 |
| Genome Sci. Tech. | Y. Zhang *et al., BMC Bioinformatics* **15**, 110 (2014). | 2 | 1 & 4 |
| Genome Sci. Tech. | M. Song *et al., Nature* **562**, 423-428 (2018). | 2 | 3 & 5 |
| Genome Sci. Tech. | M. Kleiner *et al., Nat. Commun.* **8**, 1558 (2017). | 5 | 1 & 4 |
| Genome Sci. Tech. | W. Liu *et al., Plant Biotechnol. J.* **12**, 1015-1026 (2014). | 1 | 3 & 5 |
| Microbiol. | J. Robins *et al. Vet. Res.* **46**, 68 (2015). | 3 | 1 & 4 |
| Microbiol. | M. M. McDaniel, N. Krishna, W. G. Handagama, S. Eda, V. V. Ganusov, *Front. Microbiol.* **7**, 862 (2016). | 4 | 1 & 4 |
| Microbiol. | E. Ibarguen-Mondragon, L. Esteva, E. M. Burbano-Rosero, *Math. Biosci. Eng.* **15**, 407-428 (2018). | 6 | 2 & 6 |
| Microbiol. | B. Knowles *et al., Nature* **531**, 466-470 (2016). | 5 | 2 & 6 |
| Microbiol. | S. Abel *et al., Nat. Methods* **12**, 223-226 (2015). | 2 | 2 & 6 |
| Microbiol. | L. Jackson *et al., Elife* **7**, e30134 (2018). | 2 | 1 & 4 |
| Microbiol. | T. S. Churcher *et al., PLOS Pathog.* **13**, e1006108 (2017). | 4 | 1 & 4 |
| Microbiol. | C. R. Woese, *Microbiol. Mol. Biol. Rev.* **68**, 173-186 (2004). | 5 | 1 & 4 |
| Microbiol. | F. H. Westheimer, *Science* **235**, 1173-1178 (1987). | 4 | 2 & 6 |
| Microbiol. | D. H. Parks *et al., Nat. Biotechnol.* **36**, 996-1004 (2018). | 3 | 2 & 6 |
| Microbiol. | M. Moniruzzaman *et al., Nat. Commun.* **8**, 16054 (2017). | 6 | 2 & 6 |
| Microbiol. | H. Ma *et al., Nature* **548**, 413-419 (2017). | 3 | 2 & 6 |
| Microbiol. | X. Zeng, Y. Mo, F. Xu, J. Lin, *Mol. Microbiol.* **87**, 594-608 (2013). | 6 | 2 & 6 |
| Microbiol. | M. Baym *et al., Science* **353**, 1147-1151 (2016). | 2 | 1 & 4 |
| Microbiol. | T. Jiang *et al., Int J Parasitol* **48**, 611-619 (2018). | 2 | 1 & 4 |
| Microbiol. | D. H. Parks *et al., Nat. Microbiol.* **2**, 1533-1542 (2017). | 1 | 3 & 5 |
| Microbiol. | E. A. Boyle, Y. I. Li, J. K. Pritchard, *Cell* **169**, 1177-1186 (2017). | 6 | 3 & 5 |
| Microbiol. | M. Kim, H.-S. Oh, S.-C. Park, J. Chun, *Int. J. System. Evol. Microbiol.* **64**, 346-351 (2014). | 3 | 3 & 5 |
| Microbiol. | I. Miranda *et al., mBio* **4**, e00285-13 (2013). | 2 | 3 & 5 |
| Microbiol. | A. J. Westermann *et al., Nature* **529**, 496-501 (2016). | 1 | 3 & 5 |
| Microbiol. | S. M. Hermans *et al., Appl. Environ. Microbiol.* **83**, e02826-16 (2017). | 1 | 1 & 4 |
| Microbiol. | W. Z. Stephens *et al., mBio* **6**, e01163-15 (2015). | 6 | 1 & 4 |
| ^1^Numbers identify individual evaluators. Six evaluators conducted the initial assessments of submitted papers to identify general quantitative concepts and specific skills, each assessing eight different papers. | | | |
| ^2^Evaluators were paired into teams of two, and each team selected 16 papers to discuss and rank the importance of all concepts and skills identified in the initial analysis for each paper. | | | |

**Table S1.** Papers evaluated for quantitative concepts and skills, contributing departments and programs, and associated evaluators, University of Tennessee.
